# Adaptive Transgenerational Plasticity through Priming in Response to Neighbor Density in *Mimulus platycalyx*

**DOI:** 10.1101/2025.08.26.672516

**Authors:** Arezoo Fani, John K. Kelly

## Abstract

Transgenerational plasticity (TGP) allows organisms to epigenetically alter offspring phenotypes based on their own environmental experience. TGP is adaptive if offspring perform better when their conditions match those experienced by parents. TGP is now a well-established phenomenon in plants, with current research focusing on the range of environmental causes and the mechanisms that epigenetically translate environmental experiences into phenotypic responses. This study tests neighbor density as an environmental driver of TPG, either through induction (response determined only by parental experience) or priming (response determined by the combination of parental and offspring experience). Replicate individuals of a single genotype of *Mimulus platycalyx* were grown to maturity under three conditions: No neighbors, Medium neighbor density, or High neighbor density. Offspring produced by selfing plants within each parental treatment were grown under both crowded and uncrowded conditions and measured for height, plant size traits, and flower number. Offspring growing under crowded conditions were relatively taller, had bigger leaves, and produced more flowers if their parents experienced neighbor competition than if their parents did not. In contrast, offspring grown alone performed relatively better when their parents also grew alone and did not experience competition. This pattern of performance variation, where plants did best when their environmental conditions matched those of their parents, suggests adaptive transgenerational priming in this species.

## Introduction

Phenotypic plasticity is the ability of a single genotype to express different phenotypes in response to different environments. Transgenerational plasticity (TGP) occurs when plasticity is transmitted across generations: The environmental experiences of parental plants alter the development of their offspring. It can be mediated through a variety of mechanisms, from maternal provisioning of nutrients to the transfer of small signaling RNAs, or via DNA methylation and histone modifications [1-3]. TGP can play an important role in buffering populations against environmental fluctuations, offering short-term flexibility while longer-term genetic adaptation unfolds [4]. The advantage of TGP depends largely on the predictability of environmental conditions across generations. In cases where parental environments accurately predict those of their offspring, TGP can substantially improve offspring performance [5].

TGP encompasses two distinct kinds of progeny responses. *Transgenerational induction* occurs when parental plants experience a particular stress, e.g. attack by an insect herbivore, and this induces a consistent response in progeny [6]. For instance, the progeny of attacked parents may develop elevated values for defensive traits even if they never experience attack [7]. A more subtle form of TGP has been named *transgenerational priming* [8], where parental experience does not automatically induce phenotypic changes in offspring, but simply ‘primes’ them to respond more rapidly (or to a greater degree) if these offspring experience that same stress. Induction and priming can be distinguished in the typical factorial design of TGP experiments [9]. Plants are grown and allowed to reproduce under different environmental conditions (the parental treatment). Offspring from parents grown in each parental treatment are split among distinct environmental conditions (the offspring treatment) and then measured for phenotypes and/or performance. Analyzing offspring phenotypes with a factorial ANOVA, induction is identified by a significant *direct effect* of the parental treatment. Priming is indicated by a significant interaction between parental and offspring treatments, that is when the parental effect depends on the environment experienced by the offspring.

For either induction or priming, TPG is adaptive only if offspring phenotypes are altered in the “right” direction, that which leads to increased fitness [10]. For instance, in *Campanulastrum americanum*, offspring grown under the same specific light conditions as their parents exhibited larger Leaf area and seed mass, i.e. fitness improved in matching environments [11]. Similarly, in four perennial European species tested under drought and waterlogging stress, it was matching between parents and offspring environments that lead to greater offspring biomass and reproductive output [12]. Likewise, in *Helianthemum squamatum*, offspring of drought-stressed parents had greater relative seed mass when they also experienced drought [13].

In this study, we consider varying levels of neighbor density as an environmental agent that might produce TPG. Plants often grow near neighbors, and this can have both positive and negative effects [14]. Neighboring plants can provide benefits such as buffering against wind or grazing or through increased recruitment of pollinators. However, neighbors can also trigger intense competition for light, water, and nutrients. Plants frequently show plastic responses in the presence of conspecific neighbors. In dense stands, plants that grow taller via stem elongation can have a competitive advantage especially for light [15, 16]. However, excessive elongation is not beneficial under low density conditions owing to reduced mechanical stability [10]. Plastic responses to density shape growth, structure, and reproduction. To the extent that density shows predictable pattern across generations, it may be a useful cue for TGP. For instance, parental exposure to strong competition leads to increased offspring height under similar competitive conditions in *Taraxacum brevicorniculatum* [17].

To test whether and how TGP influences offspring responses to neighbor density, we conducted a two-generation greenhouse experiment using *Mimulus platycalyx*. The parental generation was grown under three treatments: Control (no neighbors), Medium Density, and High Density. These parental plants were self-fertilized, and seeds were collected to be used as the offspring generation. We grew from each parental treatment under both crowded and uncrowded conditions and measured them for traits related to growth and reproduction. We find no evidence for transgenerational induction but a strong signal of transgenerational priming.

## Materials and methods

### Study species

*Mimulus platycalyx* is an annual species that occupies wet habitats [18]. Compared to its close relative *Mimulus guttatus*, which occurs in a broad range of moist environments, *M. platycalyx* has a more restricted distribution [18]. It also differs morphologically, displaying a broader calyx and more upright growth than *M. guttatus. M. platycalyx* reproduces often by self-fertilization with field estimates for the outcrossing rate ranging from 0.1 to 0.5 [19-21]. These estimates are upwardly biased owing to inbreeding depression, and thus, *M. platycalyx* is likely to be predominantly selfing in most populations in most years [22]. Local populations of *M. platycalyx* do exhibit genetic variation in quantitative traits, although levels appear to be quite reduced relative to *M. guttatus* [22, 23]. The mating system of *M. platycalyx* and its ecological and morphological characteristics make it a compelling species for studying TGP. Limited genetic variation within *M. platycalyx* populations may restrict their potential for rapid genetic evolution. In such cases, TGP may provide a mechanism for coping with recurring environmental conditions.

The seeds of *Mimulus platycalyx* used in this experiment were derived from a collection by Noland Martin from a population near Guenoc Winery, Napa County, California (38.756913, -122.607345; population code “NAP”). A single inbred line was established through successive generations of self-pollination in the greenhouse. This line was selfed for at least five generations prior to the present experiment. All plants used in this experiment, both parental and offspring generations, were sampled from the progeny of a single family within this inbred line. These plants are highly homozygous and should be (nearly) equivalent in genotype.

### Experimental Design

We grew 93 genetically identical plants (the parents), between 29 and 32 in each of three distinct treatments (Figure 1). Control parents grew alone in a pot, experiencing no neighbor competition. Medium Density parents were surrounded by 8 neighbors of the same genotype at the edge of the pot, sufficient to generate moderate competition. High Density parents were surrounded by a dozen very close neighbors in addition to 8 “edge” neighbors. Pilot experiments had shown the High Density condition induced a pronounced within generation plastic response of the focal plant involving stem elongation. All the neighbors were the same genotype as the focal plants. After five weeks, close neighbors were removed in the High density pots to allow for focal plant growth (edge neighbors remained). To ensure sufficient seed production, the parental plants were hand self-pollinated. We collected, counted, and weighed the total set of seeds produced by each parental generation plant. “Mean seed mass”, as reported below, is the total mass divided by seed number for each parental plant.

**Figure 1.**
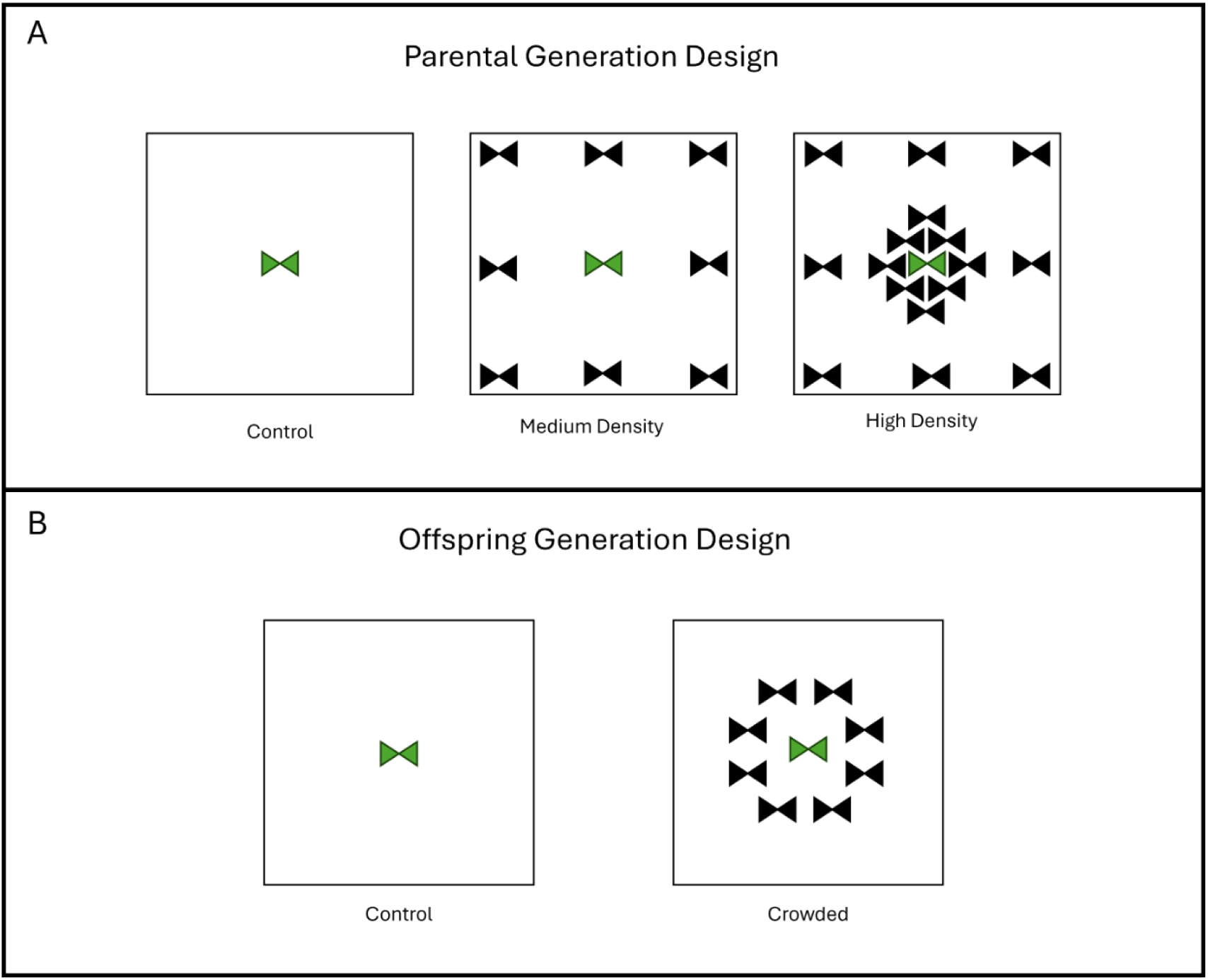
**The treatments are depicted for the (A) Parental and (B) Offspring generations. Focal plants are shown in green, and neighbor plants are shown in black.**

The offspring generation was initiated three weeks after the completion of seed collection from the parental generation. Offspring generation seeds were stratified for the last 7 days of this interval on wet soil at 4°C. After germination, offspring within maternal families were randomly split between two growth treatments (figure 1, lower panel). In “Control”, focal plants were not surrounded by any neighbors. In “Crowded”, focal plants were surrounded by eight close neighbors, forming a ring around it. In terms of shading, the Crowded offspring treatment is intermediate to the Medium and High Density treatments of the parental generation. It was not practical to repeat the High Density treatment in the offspring generation because we could not identify and track a single focal plant within known ancestry. Offspring pots were assigned to ‘blocks,’ defined by a shared start date (pots were initiated in two cohorts staggered by one week) and grouped together within the same flat. Although the members of a block were spatially separated from other offspring pots, they were located in same area of greenhouse at any specific time. Flats were rotated twice per week. Importantly, the experimental treatments (all combinations of parental and offspring treatment) were balanced across blocks. This means that blocks effects can be statistically factored out when testing for treatment effects.

Plants were grown under standard greenhouse conditions for *Mimulus* [24], with an 16-hour photoperiod, daytime temperature of 24°C, and nighttime temperature of 15°C. Berger MB6 All Purpose Mixed Soil was used with a fungicide (Subdue GR, Syngenta) and a pesticide (Marathon 1%, OHP) added at 20 ml and 250 ml per 25 gallons of soil, respectively. All plants were grown in 4-inch pots filled with the standardized soil mixture. All plants received 10N–30P–20K Blossom Booster fertilizer (Jack’s Classic) applied weekly at a concentration of 250 ml per 25 gallons of water. During early germination and growth, plants were top misted daily and subsequently received regular bottom watering for the duration of the experiment.

### Data Collection & Measured Traits

Measurements were taken over the course of the experiment, but here we report and analyze only those from the final census taken ca. 42 days after seed-to-soil. At harvest, leaf width was measured on the largest leaf. This measure of leaf size is strongly associated with the total above ground biomass of plants in *Mimulus* [25]. All internodal lengths taken along with total plant height. The length of the pistil was measured on two flowers given that previous studies have shown this to be a stronger predictor of the maximum seed set of a flower (seed set of flowers when they are given a saturating dose of pollen; [26]). Finally, we also recorded the total number of flowers produced by each plant up to the final harvest. Dimension measurements were taken using a 6-inch stainless steel drafting ruler (General Tools & Instruments, USA) to 1/100^th^ inch precision.

### Statistical Analysis

We used general linear models (ANOVAs and ANCOVAs) to test treatment effects on all measured traits. For average seed mass of parental plants, we applied a single factor ANOVA with Parental treatment (fixed) as the factor. For final offspring values of height, internodal differences, leaf size, pistil length, and flower number, we first fit a factorial ANOVA with Parental Treatment and Offspring Treatment (both fixed) and their interaction. This model also included block and parental line (nested within Parental Treatment) as random effects. Finally, we reran the model fits for each offspring trait including the average seed mass of each parental line as a covariate. Mean seed mass was first used as a simple linear predictor and then we allowed it to interact with Offspring treatment in these ANCOVA tests. All statistical analyses were performed using Minitab (version 21.2, 64-bit).

## Results

A total of 587 offspring plants were measured at the final harvest. For each of the four composite measures (final height, flower number, maximum leaf width, and mean pistil length), there was a highly significant negative effect of the offspring Crowded treatment (Figure 2, Table 1). This indicates that neighbor interactions were competitive.

**Table 1.**
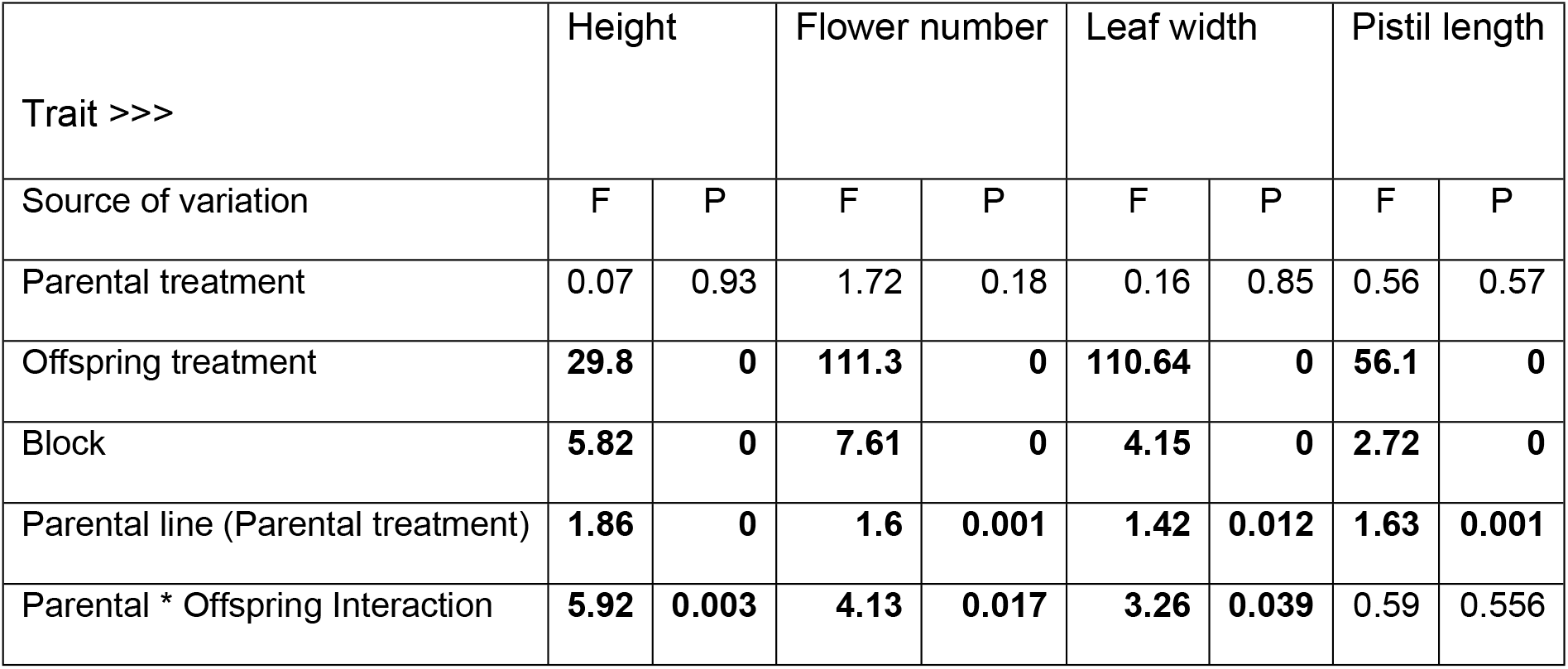
**Results of factorial ANOVA testing the effects of parental treatment, offspring treatment, and their interaction are presented for four offspring traits. F ratios and associated p-values are reported for each factor. Significant effects (p < 0.05) are shown in bold; values of 0 indicate p < 0.0005.**

**Figure 2.**
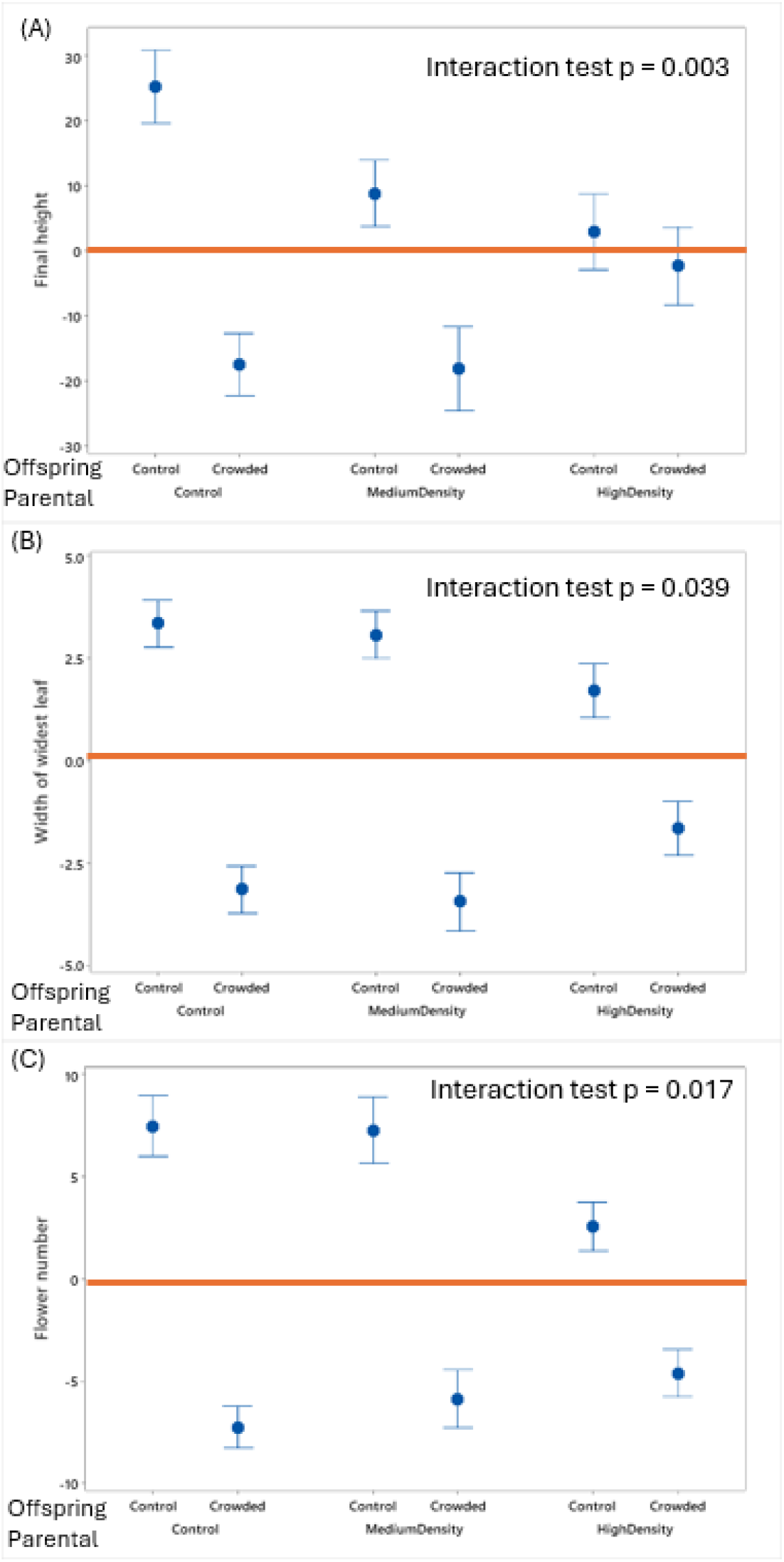
**The mean values for (A) final height, (B) leaf width, and (C) flower number are reported for each combination of parental and offspring treatment. Values are reported relative the overall mean value for each trait (orange bar): 231mm for height, 28.0mm for leaf, and 23.56 for flower number. The p-value from the Parental x Offspring treatment interaction is reported in each panel. Error bars are +/-1 SEM after factoring out the effects of block.**

Parental treatment had no direct effect on any trait. However, three of the four traits exhibited a significant Parent by Offspring treatment interaction (Table 1). Only average pistil length was unresponsive to parental treatment. For the other three traits, the difference between Crowded and Control offspring changed depending on the Parental treatment (Figure 2). For all offspring traits, parental line explained a significant portion of the variation (Table 1).

In the Offspring Control treatment, plants were taller when their parents had been grown alone (Parental Control) compared to when their parents had experienced High Density conditions (average difference ca. 22mm). In contrast, for offspring grown in the Crowded Treatment, they finished substantially shorter if their parents were grown alone relative to parents grown under High Density (average difference ca. -15mm). For flower number and leaf width, the same direction of differences obtained. Control offspring had more flowers (average difference = 4.9) and larger leaves (average difference = 1.65mm) if their parents were grown without neighbors. Offspring in the Crowded treatment had fewer flowers (average difference = -2.6) and smaller leaves (average difference = -1.51mm) if their parents were grown without neighbors. Responses to the Medium Density parent treatment were generally intermediate, although perhaps more like the Control than High Density parental treatment (Figure 2).

The plastic growth responses of offspring are assessed by the lengths of different internodal segments (Figure 3). Regardless of parental treatment, internodal lengths tended to increase through time when offspring grew alone (green arrows in Figure 3) and decline when offspring experienced crowding (orange arrows). This is expected if shading increases with time in the Crowded Treatment but not the offspring Control Treatment. Shading should reduce the *relative* growth rate in the Crowded Treatment. However, the origins and lengths of the arrows in Figure 3 reveal the nature of Parental treatment effects. Considering offspring grown without neighbors, the first internode was about the same regardless of parental experience. However, offspring of Control Parents grew much more rapidly than offspring of High Density parents when the offspring were grown without neighbors. In contrast, the early growth (expansion of the first internode) differed substantially among parental treatments when offspring were grown in the Crowded treatment. This suggests an ‘anticipatory’ effect: Offspring of parents that experienced either Medium or High Density get a head start when growing with neighbors.

**Figure 3.**
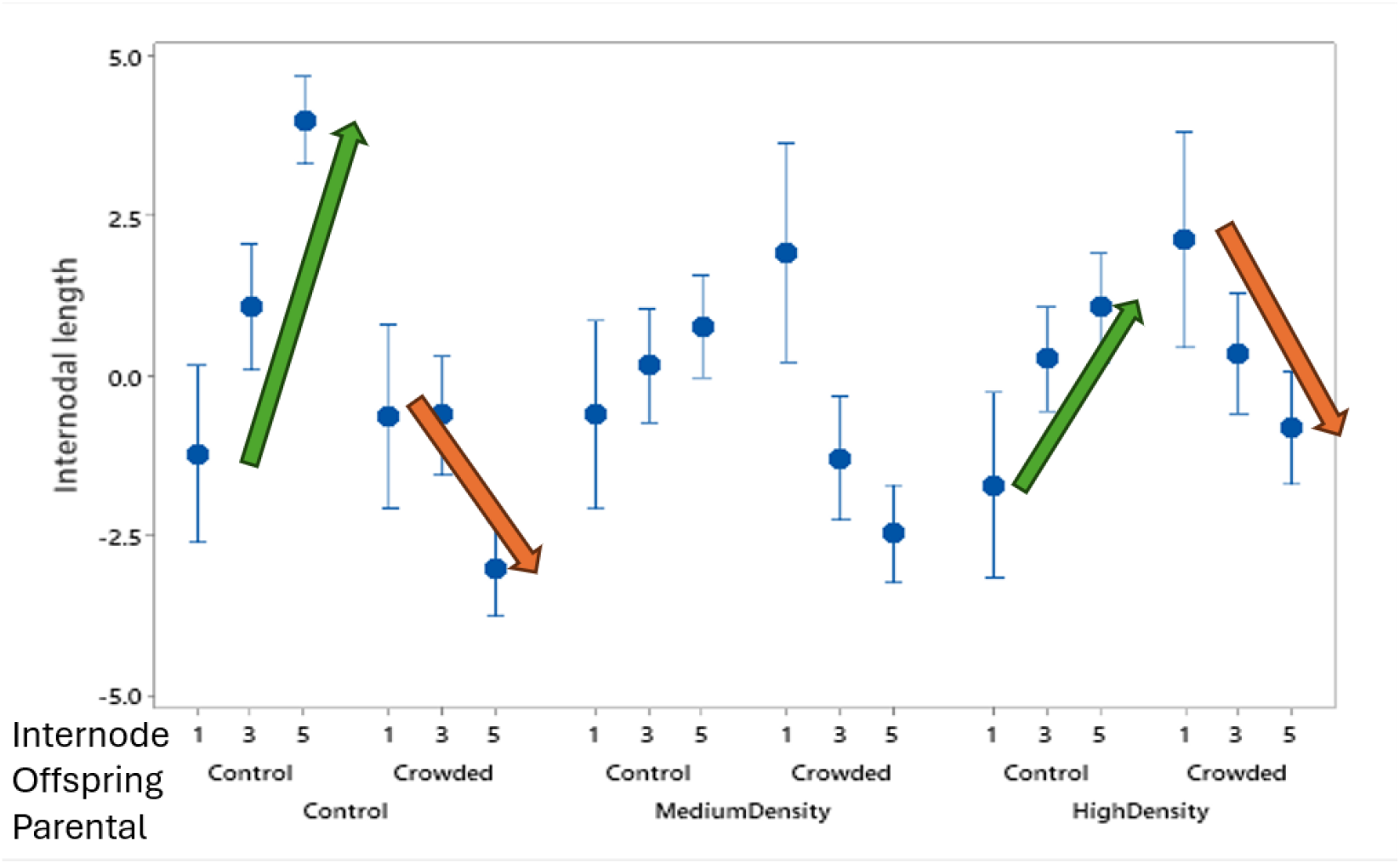
**The average internodal distances are reported for the 1^st^, 3^rd^, and 5^th^ internodes for each combination of parental and offspring. Green and orange arrows highlight the change in internode length across developmental time, with green indicating growth in control conditions and orange in crowded conditions. As in Figure 2, values are standardized relative to the mean length of the relevant internode and error bars are +/-1 SEM.**

Parental treatment did influence the average mass of seeds (Figure 4A). Seeds from plants in the Medium and High Density conditions were significantly heavier than those produced by Control parents (F_2,90_ = 9.15, p < 0.0005). To evaluate seed mass effects on offspring generation responses, we determined the mean phenotype of offspring from each parental family (we do not have the mass of the individual seeds that produced specific offspring). There was no relationship between seed mass and average offspring phenotype of each family (all regressions non-significant, Supplemental Figure 1). However, an effect of seed mass becomes apparent if we split the offspring in each family according to the treatment that offspring experienced. For height, there was a positive effect of seed mass among crowded offspring but not control offspring, the test for slope heterogeneity is highly significant (F_1,176_ = 11.54, p = 0.001). Considering each offspring treatment separately, the regression is significantly positive in the Crowded treatment but non-significant in the Control. Also, if one subdivides these groups based on Parental treatment, there is no evidence of slope heterogeneity based on Parental treatment. This analysis was applied to the other three traits (Supplemental Figure 2). Slope heterogeneity based on offspring treatment was significant for leaf width (F_1,176_ = 8.58, p = 0.004), but not for flower number (F_1,176_ = 2.51, p > 0.05) or pistil length (F_1,176_ = 0.01, p > 0.05).

**Figure 4.**
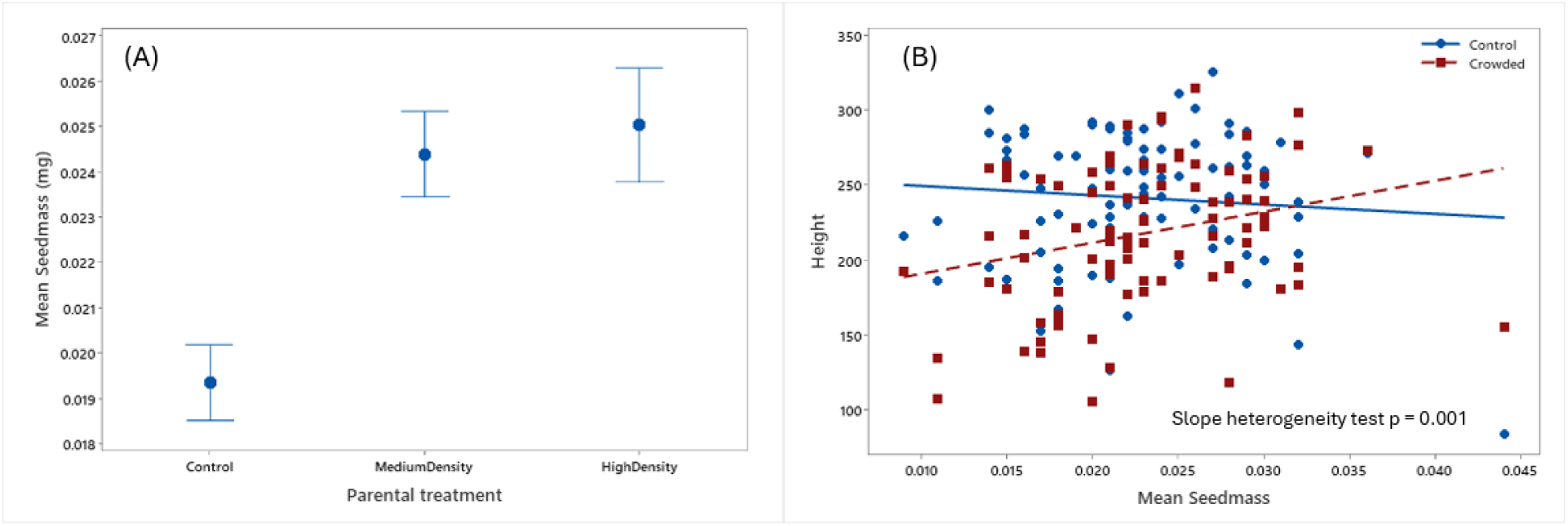
**(A) The mean seed mass produced by parental plants is reported for each treatment group. (B) The regressions of family mean offspring height for either the Control (blue) or Crowded (red) offspring treatment onto mean seed mass is reported with the statistical test for slope heterogeneity.**

Noting these dependencies, we expanded the ANOVA model (Table 1) to include covariates based on mean parental seed mass. This ANCOVA, reported as Table 2, includes two additional terms (“Seed mass” and “Seed mass * Offspring treatment”) that characterize seed mass effects on offspring phenotypes in a way that is specific to offspring treatment. As expected from Figure 4B, we find a highly significant effect of Seed mass * Offspring treatment for Height and Leaf width. The test on factors in this ANCOVA evaluate the effects of parental and offspring treatments after statistically removing the effect of mean parental seed mass. Importantly, the interactions between parental and offspring treatments, which were significant for height, flower number and leaf width in the ANOVAs (Figure 2, Table 1), become marginally non-significant in the ANCOVAs (Table 2).

**Table 2.**
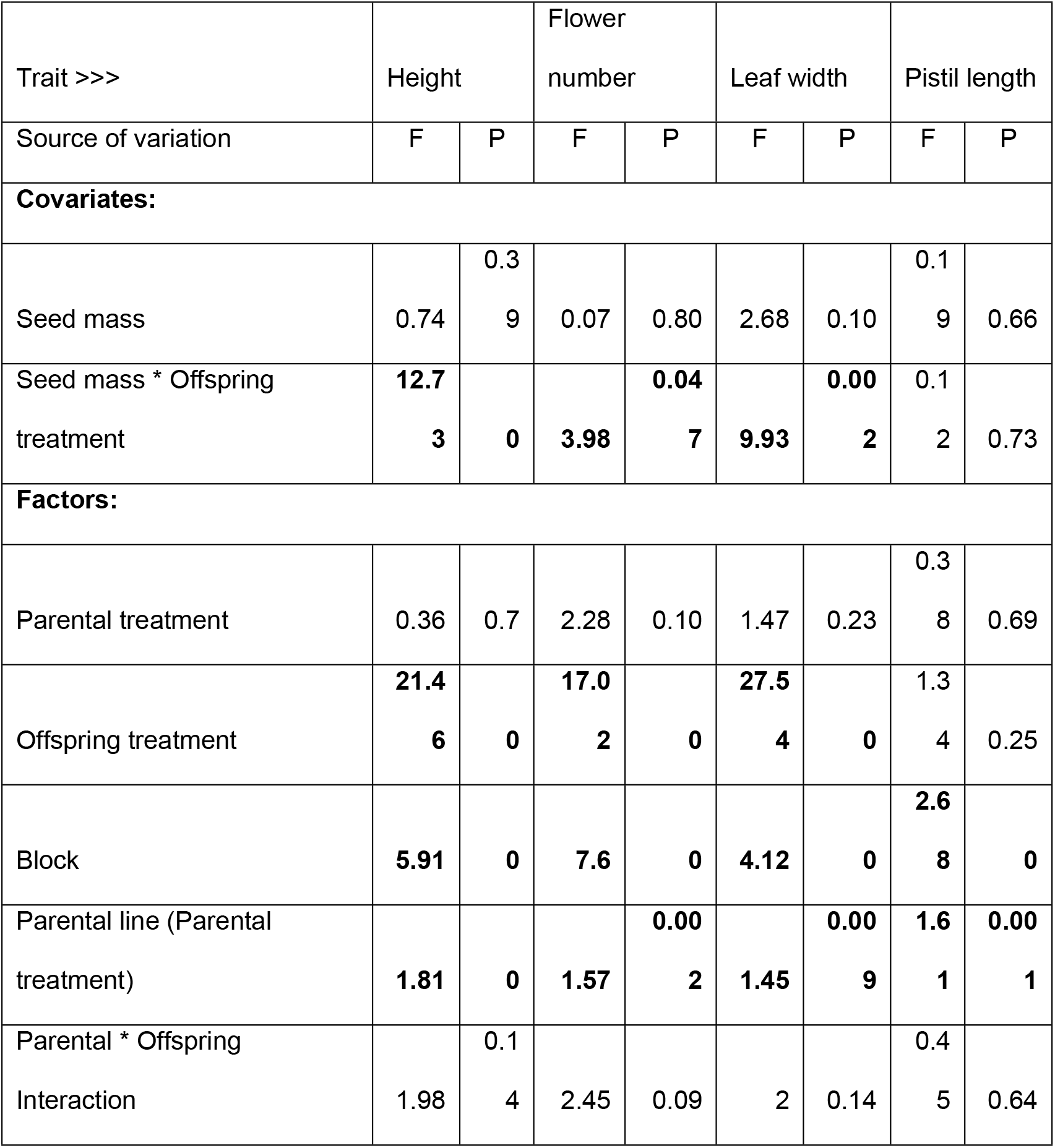
**Results of ANCOVA testing the effects of parental treatment, offspring treatment, their interaction, and covariates (seed mass and seed mass × offspring treatment) on four offspring traits. F ratios and associated p-values are reported for each factor. Significant effects (p < 0.05) are shown in bold; values of 0 indicate p < 0.0005.**

## Discussion

This study demonstrates transgenerational plasticity (TGP) in *Mimulus platycalyx* in response to varying levels of competition from conspecific neighbors. The results favor transgenerational priming rather than transgenerational induction. The offspring of parents that experienced competition exhibit phenotypic changes that appear to increase their competitive ability, but only if these offspring themselves experience crowding (Figure 2). Because flower size (and the estimated reproductive capacity per flower) was unaffected by parental treatment, we can use the total flower numbers as an integrated performance measure for this experiment. Progeny growing without neighbors produced 19% *more* flowers if their own parents had not experienced any competition (parental Control) than if they had experienced intense shading (parental High Density). Progeny growing under crowded conditions produced 14% *fewer* flowers if their own parents had not experienced any competition than if they were in the High Density treatment. This reversal of relative fitness with environment is the signature of adaptive plasticity [10].

Our experiment had three distinct treatments corresponding to different levels of intraspecific competition (Figure 1). Generally, parents that experienced moderate competition produced a more modest progeny response than those experiencing high competition. This suggests that the quantitative level of stress experienced by parents can regulate the magnitude of the priming response with more intense conditions producing stronger transgenerational effects [27]. This may not be surprising given that the proximate drivers of TGP (nutrients, signaling molecules, levels and patterns of methylation and histone modifications) all vary quantitatively.

Across the traits that we measured, the signal of priming was strongest for final plant height (Figure 2A). Height is a key trait when plants are competing for light under high density conditions. Taller plants often gain competitive advantages in dense stands by overtopping neighbors to capture light [16], but excessive stem elongation can be detrimental when neighbors are not present. Final height is as much an outcome as a growth strategy; it being determined by growth/allocation decisions early in the lifetime of a plant (Figure 3). A potential cost of priming is suggested by the green arrows in Figure 3: Offspring of parents that experienced competition showed reduced *relative* growth when they grew without competitors. In contrast, primed offspring got going more quickly in the presence of competitors (orange arrows in Figure 3) than offspring from the Control parental treatment. Puy, de Bello (17) obtained similar results in a recent experiment examining TPG in response to competition in *Taraxacum brevicorniculatum*.

### To prime or to induce?

The closest relative to *M. platycalyx* that has been shown to exhibit TPG is *M. guttatus*, which shows a very strong induction response to both natural and simulated herbivory. The response is not limited to growth and morphological features of offspring, but also gene expression, genomewide DNA methylation patterns, and herbivory resistance in the field [7, 28-31]. The present experiment of *M. platycalyx* shows no evidence of induction – the direct effect of parental treatment is never even close to significant (Table 1). Why should *M. platycalyx* prime their offspring for neighbor competition when *M. guttatus* induces defense for herbivore attack? Whether to prime or induce may be determined by developmental or physiological constraints. Priming requires a quicker developmental response than induction because offspring wait to receive the environmental stimulus before altering gene expression, hormone production, etc., to produce phenotypic changes. Neighbor competition might be a good candidate for rapid response given that auxin mediated stem elongation can occur within minutes to few hours upon shading [32, 33]. The defense phenotypes documented in *M. guttatus* (where TGP occurs by induction), such as the development of glandular trichomes and production of defensive secondary compounds, cannot be produced as rapidly. Consequently, plants may be devoured before they can mount an effective defense by priming.

Ecological circumstances may also determine whether priming or induction is favorable [34]. Some degree of predictability of environmental conditions from the parental to offspring generation is necessary for any form of TGP to be adaptive. In “climax” populations of *M. platycalyx*, where the species exists in high density (nearly) monospecific stands, parental experience of density should be reliable. However, like other annual herbaceous species, density may decline substantially in years where idiosyncratic environmental perturbations cause low germination or high offspring mortality. Also, persistence of the *M. platycalyx* metapopulation requires the occasional colonization event of newly available habitat patches. A seed produced from a high density population that establishes in a new area will certainly experience a different competitive environment than its parents. In such circumstances, it would be advantageous to sense the environment before committing to a phenotype based on parental experience.

### The subtle effect of seed mass and the basis of transgenerational priming

Parental plants that experienced competition produced heavier seeds (Figure 4A). While stress often reduces seed provisioning in plants [9], a number cases like *M. platycalyx* have been documented [35]. For example, drought stress causes plants to increase seed mass in *Helianthemum squamatum* [13]. A very interesting result is that if we add seed mass as a simple covariate when analyzing offspring phenotypes, it has no effect on any trait (Supplemental Figure S1). However, when we allow seed mass to have differing effects for control versus crowded offspring, a highly significant pattern emerges. Both height and flower number increase with mean seed mass, but only if the offspring experience crowding (Figure 4B, Supplemental Figure S2). Since priming is indicated by an interaction between parental and offspring treatments, Table 2 suggests that priming is at least partially mediated through seed mass or some other (unmeasured) factor that is correlated with seed mass.

The prediction of priming from seed mass does not imply that simple nutrient provisioning explains TGP. If increased seed size translates directly to increased early growth, we should have seen positive effects in both control and crowded conditions. For offspring grown without neighbors, seed mass actually had weak negative correlations (non-significant) with final size traits (Supplemental Figure S2). The conditionality of the response suggests that other epigenetic factors may be involved. While factors such as DNA methylation have not yet been studied in *M. platycalyx*, they have been implicated in TGP of the closely related *M. guttatus* [30]. In *M. guttatus*, the epigenetic signal of TGP for trichomes is transmitted as much thorough male gametes as female gametes [31], which clearly requires an explanation beyond seed nutrient provisioning. More work is needed to determine the proximate mechanisms of priming in *M. platycalyx*.

## Conclusion

Our findings demonstrate that parental exposure to competition from neighboring plants leads to transgenerational priming in *Mimulus platycalyx*. Significant Parent × Offspring treatment interactions for height, leaf width, and flower number (Table 1, Figure 2) indicate that offspring responses depend on both their own environment and that of their parents. Analyses of internode growth (Figure 3) inform questions on the developmental basis of anticipatory plasticity, with offspring of crowded parents performing better under crowding but relatively worse when growing alone. Parental treatment also altered seed mass (Figure 4A), with crowding having the counter-intuitive effect of increasing average seed size. The size of seed had no average effect on offspring traits. However, it had a contingent effect by increasing particular traits but only when offspring experienced crowding; a result that provides valuable clues about the mechanistic basis of priming. These results highlight the value of priming relative to induction as a form of TPG because it provides plants additional flexibility in plastic responses to match current environmental conditions.

## Data availability

All data used for this paper have been archived in Dryad (DOI pending).

## Supplemental Figures

**Figure S1.**
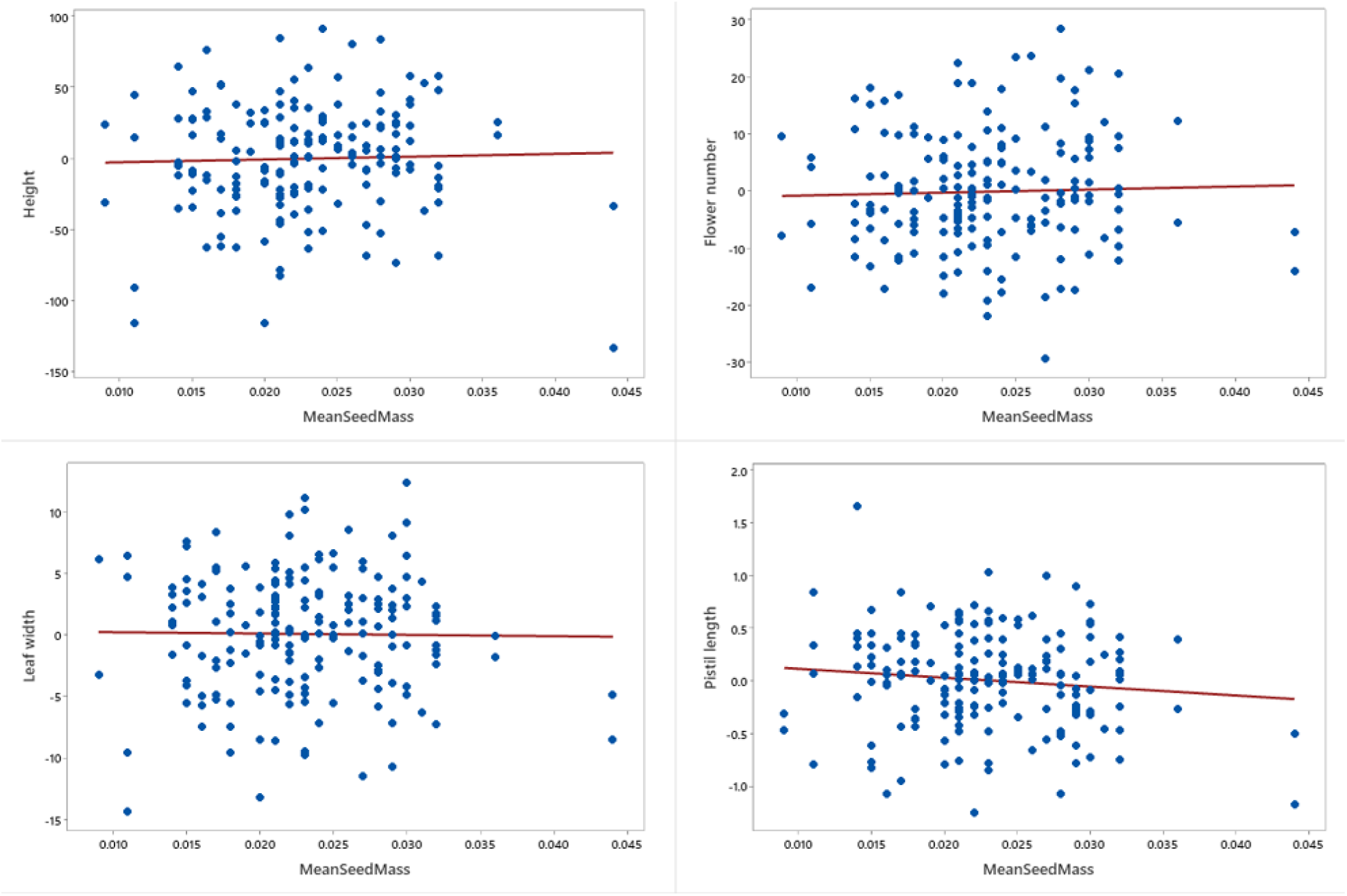
**The mean offspring trait value per family is reported as a function of mean parental seed mass for that family. Each point represents a family, and red lines show the linear regression fit. None of the slopes are significantly different from zero (p > 0.05), indicating no relationship between parental seed mass and offspring trait expression across families.**

**Figure S2.**
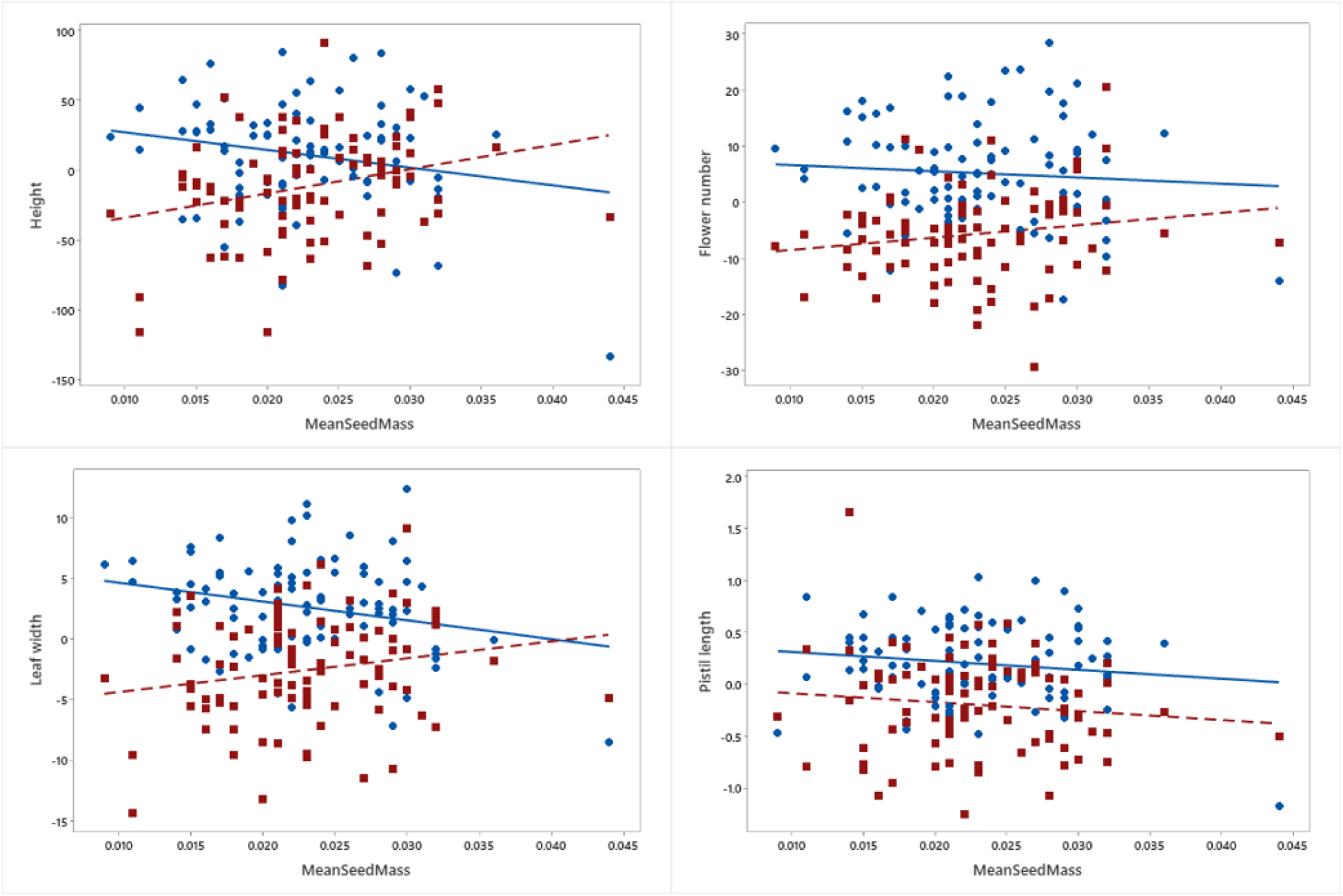
**Regressions of mean offspring trait values on mean parental seed mass are shown separately for Control (blue) and Crowded (red) offspring treatments. Each point represents a family, and lines indicate fitted linear regression models.**

